# The Patchy Distribution of Restriction-Modification System Genes and the Conservation of Orphan Methyltransferases in Halobacteria

**DOI:** 10.1101/551721

**Authors:** Matthew S. Fullmer, Matthew Ouellette, Artemis S. Louyakis, R. Thane Papke, J. Peter Gogarten

**Affiliations:** Department of Molecular and Cell Biology, University of Connecticut, Storrs, CT, USA; Bioinformatics Institute, School of Biological Sciences, The University of Auckland, Auckland, New Zealand

**Keywords:** HGT, Restriction, Methylation, Gene Transfer, Selfish Genes, Archaea, Haloarchaea, DNA methylase, epigenetics

## Abstract

Restriction-modification (RM) systems in Bacteria are implicated in multiple biological roles ranging from defense against parasitic genetic elements, to selfish addiction cassettes, and barriers to gene transfer and lineage homogenization. In Bacteria, DNA-methylation without cognate restriction also plays important roles in DNA replication, mismatch repair, protein expression, and in biasing DNA uptake. Little is known about archaeal RM systems and DNA methylation. To elucidate further understanding for the role of RM systems and DNA methylation in Archaea, we undertook a survey of the presence of RM system genes and related genes, including orphan DNA methylases, in the halophilic archaeal class Halobacteria. Our results reveal that some orphan DNA methyltransferase genes were highly conserved among lineages indicating an important functional constraint, whereas RM systems demonstrated patchy patterns of presence and absence. This irregular distribution is due to frequent horizontal gene transfer and gene loss, a finding suggesting that the evolution and life cycle of RM systems may be best described as that of a selfish genetic element. A putative target motif (CTAG) of one of the orphan methylases was underrepresented in all of the analyzed genomes, whereas another motif (GATC) was overrepresented in most of the haloarchaeal genomes, particularly in those that encoded the cognate orphan methylase.

## 1. Introduction

DNA methyltransferases (MTases) are enzymes which catalyze the addition of a methyl group to a nucleotide base in a DNA molecule. These enzymes will methylate either adenine, producing *N*6-methyladenine (6mA), or cytosine, producing either *N*4-methylcytosine (4mC) or C5-methylcytosine (5mC), depending on the type of MTase enzyme [1]. DNA methyltransferases typically consist of three types of protein domains: an S-adenosyl-L-methionine (AdoMet) binding domain which obtains the methyl group from the co-factor AdoMet, a target recognition domain (TRD) which binds the enzyme to the DNA strand at a short nucleotide sequence known as the recognition sequence, and a catalytic domain which transfers the methyl group from AdoMet to a nucleotide at the recognition sequence [2]. The order in which these domains occur in a MTase varies and can be used to classify the enzymes into the subtypes of *α, β, γ, δ, ε*, and *ζ* MTases [3–5].

In bacteria and archaea, MTases are often components of restriction-modification (RM) systems, in which an MTase works alongside a cognate restriction endonuclease (REase) that targets the same recognition site. The REase will cleave the recognition site when it is unmethylated, but the DNA will escape cutting when the site has been methylated by the MTase; this provides a self-recognition system to the host where it differentiates between its own methylated DNA and that of unmethylated, potentially harmful foreign DNA that is then digested by the host’s REase [6–8]. RM systems have also been described as addiction cassettes akin to toxin-antitoxin systems, in which post-segregational killing occurs when the RM system is lost since the MTase activity degrades more quickly than REase activity, resulting in digestion of the host genome at unmodified recognition sites [9,10]. RM systems have been hypothesized to act as barriers to genetic exchange and drive population diversification [11,12]. In *Escherichia coli*, for example, conjugational uptake of plasmids is reduced by the RM system EcoKI when the plasmids contain EcoKI recognition sequences [13]. However, transferred DNA that is digested by a cell’s restriction endonuclease can still effectively recombine with the recipient’s chromosomal DNA [7,14,15]; the effect of DNA digestion serves to limit homologous recombinant DNA fragment size [16]. Restriction thus advantages its host by decreasing transfer of large mobile genetic elements and infection with phage originating in organisms without the cognate MTase [8], while also reducing linkage between beneficial and slightly deleterious mutations [17].

There are four major types of RM systems which have been classified in bacteria and archaea [18,19]. Type I RM systems consist of three types of subunits: REase (R) subunits, MTase (M) subunits, and site specificity (S) subunits which contain two tandem TRDs. These subunits form pentamer complexes of two R subunits, two M subunits, and one S subunit, and these complexes will either fully methylate recognition sites which are modified on only one DNA strand (hemimethylated) or cleave the DNA several bases upstream or downstream of recognition sites which are unmethylated on both strands [20,21]. The MTases and REases of Type II RM systems have their own TRDs and operate independently of each other, but each one targets the same recognition site [22]. There are many different subclasses of Type II RM system enzymes, such as Type IIG enzymes which contain both REase and MTase domains and are therefore capable of both methylation and endonuclease activity [23]. Type III RM systems consist of REase (Res) and MTase (Mod) subunits which work together as complexes, with the Mod subunit containing the TRD which recognizes asymmetric target sequences [24]. Type IV RM systems are made up of only REases, but unlike in other RM systems, these REases will target and cleave methylated recognition sites [20,25].

MTases can also exist in bacterial and archaeal hosts as orphan MTases, in which they occur independently of cognate restriction enzymes and typically have important physiological functions [26]. In *E. coli*, the orphan MTase Dam, an adenine MTase which targets the recognition sequence GATC, is involved in regulating the timing of DNA replication by methylating the GATC sites present at the origin of replication (*oriC*) [27]. The protein SeqA binds to hemimethylated GATC sites at *oriC*, which prevents re-initiation of DNA replication at *oriC* after a new strand has been synthesized [28,29]. Dam methylation is also important in DNA repair in *E. coli*, where the methylation state of GATC sites is used by the methyl-directed mismatch repair (MMR) system to identify the original DNA strand in order to make repairs to the newly-synthesized strand [30–32]. In *Cauldobacter crescentus*, the methylation of target sites in genes such as *ctrA* by orphan adenine MTase CcrM helps regulate the cell cycle of the organism [33–35]. The importance of orphan MTases in cellular processes is likely the reason why they are more widespread and conserved in bacteria compared to MTases associated with RM systems [36,37].

MTases and RM systems have been well-studied in the bacteria, but less research has been performed in archaea, with most studies focused on characterizing RM systems of thermophilic species [38–42]. Recent research into the halophilic archaeal species *Haloferax volcanii* has demonstrated a role for DNA methylation in DNA metabolism, and probably uptake: cells could not grow on wild type *E. coli* DNA as a phosphorous source, whereas unmethylated *E. coli* was metabolized completely [43,44]. In an effort to better understand this phenomenon, we characterized the genomic methylation patterns (methylome) and MTases in the halophilic archaeal species *Haloferax volcanii* [45,46]. However, the distribution of RM systems and MTases among the archaea has not been extensively studied, and thus their life histories and impact on host evolution are unclear.

To that end we surveyed the breadth of available genomes from public databases representing the class Halobacteria, also known as the Haloarchaea, for RM system and MTase candidate genes. We further sequenced additional genomes from the genus *Halorubrum* which provided an opportunity to examine patterns among very closely related strains. Upon examining their patterns of occurrence, we discovered orphan methyltransferases widely distributed throughout the Haloarchaea. In contrast, RM system candidate genes had a sparse and spotty distribution indicating frequent gene transfer and loss. Even individuals from the same species isolated from the same environment and at the same time, differed in the RM system complement.

## 2. Materials and Methods

### Search Approach

The starting data consists of 217 Halobacteria genomes from NCBI and 14 in-house sequenced genomes (Supplementary Table S1). We note that some of these genomes were assembled from shotgun metagenome sequences and not from individual cultured strains. Genome completion was determined through identification of 371 Halobacteriaceae marker genes using CheckM v1.0.7 [47]. Queries for all restriction-methylation-specificity genes were obtained from the Restriction Enzyme dataBASE (REBASE) website [48,49]. As methylation genes are classified by function rather than by homology [48] the protein sequences of each category were clustered into homologous groups (HGs) via the *uclust* function of the USEARCH v9.0.2132 package [50] at a 40 percent identity. The resulting ∼36,000 HGs were aligned with MUSCLE v3.8.31 [51]. HMMs were then generated from the alignments using the *hmmbuild* function of HMMER3 v3.1b2 (hmmer.org). The ORFs of the 217 genomes were searched against the profiles via the *hmmsearch* function of HMMER3. Top hits were extracted and cross hits filtered with in-house Perl scripts. Steps were taken to collapse and filter HGs. First, the hits were searched against the arCOG database [52] using BLAST [53] to assign arCOG identifiers to the members of each group. Second the R package *igraph* v1.2.2 [54] was used to create a list of connected components from the arCOG identifications. All members of a connected component were collapsed into a single collapsed HG (cHG).

Because REBASE is a database of all methylation-restriction-related activities there are many members of the database outside our interest. At this point we made a manual curation of our cHGs attempting to identify known functions that did not apply to our area of interest. Examples include protein methylation enzymes, exonucleases, cell-division proteins, etc. The final tally of this clustering and filtering yielded 1696 hits across 48 total candidate cHGs. arCOG annotations indicate DNA methylase activity, restriction enzyme activity, or specificity module activity as part of an RM system for 26 cHGs. The remaining 22 cHGs had predominant arCOG annotations matching other functions that may reasonably be excluded from conservative RM system-specific analyses. For a graphical representation of the search strategy see supplementary materials **Figure S1**. The putative Type IV methyl-directed restriction enzyme gene, *mrr*, which is known to be present in *Haloferax volcanii* had not been assembled into an cHG. We assembled a cluster of *mrr* homologs and determined their presence and absence using Mrr from *Haloferax volcanii* DS2 (accession: ADE02322.1) as query in BLASTP searches against each genome (E-value cut-off 10^−10^).

### Reference Phylogeny

A reference tree was created using the full complement of ribosomal proteins. The ribosomal protein set for *Halorubrum lacusprofundi* ATCC 49239 was obtained from the BioCyc website [55]. Each protein orf was used as the query in a BLAST [53] search against each genome. Hits for each gene were aligned with MUSCLE v3.8.31 [51] and then concatenated with in-house scripting. The concatenated alignment was subjected to maximum likelihood phylogenetic inference in the IQ-TREE v1.6.1 suite with ultrafast bootstrapping and automated model selection [56,57]. The final model selection was LG+F+R9.

### F81 Presence-Absence Phylogeny

It is desirable to use maximum-likelihood methodology rather than simple distance measures. To realize this, the matrix was converted to an A/T alignment by replacing each present with an “A” and absent with a “T.” This allowed use of an F81 model with empirical base frequencies. This confines the base parameters to only A and T while allowing all of the other advantages of an ML approach. IQ-TREE was employed to infer the tree with 100 bootstraps [57].

### Horizontal Gene Transfer Detection

Gene trees for each of the cHGs were inferred using RAxML v8.2.11 [58] under PROTCATLG models with 100 bootstraps. The gene trees were then improved by resolving their poorly supported in nodes to match the species tree using TreeFix-DTL [59]. Optimized gene tree rootings were inferred with the OptRoot function of Ranger-DTL. Reconciliation costs for each gene tree were computed against the reference tree using Ranger-DTL 2.0 (http://compbio.engr.uconn.edu/software/RANGER-DTL/) [60] with default DTL costs. One-hundred reconciliations, each using a different random seed, were calculated for each cHG. After aggregating these with the AggregateRanger function of Ranger-DTL the results were summarized and each prediction and any transfer inferred in 51% or greater of cases was counted as a transfer event.

### Data Analysis and Presentation

The presence-absence matrix of cHGs was plotted as a heatmap onto the reference phylogeny using the *gheatmap* function of the R Bioconductor package *ggtree* v1.14.4 [61,62]. The rarefaction curve was generated with the *specaccum* function of the *vegan* v2.5-3 package in R [63] and number of genomes per homologous group was plotted with ggplot2 v3.1.0 [64]. Spearman correlations and significances between the presence-absence of cHGs was calculated with the *rcorr* function of the *hmisc* v4.1-1 package in R (http://biostat.mc.vanderbilt.edu/wiki/Main/Hmisc). A significance cutoff of p < 0.05 was used with a Bonferroni correction. All comparisons failing this criterion were set to correlation = 0. These data were plotted into a correlogram via the *corrplot* function of the R package *corrplot* v0.84. To compare the Phylogeny calculated from Presence-Absence data to the ribosomal protein reference, the bootstrap support set of the presence-absence phylogeny was mapped onto the ribosomal protein reference tree using the *plotBS* function in *phangorn* v2.4.0 [65]. Support values equal to or greater than 10% are displayed. To compare phylogenies using Internode Certainty, scores were calculated using the IC/TC score calculation algorithm implemented in RAxML v8.2.11 [58,66].

### Synteny

Genomes were searched for location of cHGs. Proximity was used to determine synteny of groups of cHGs frequently identified on the same genomes.

### Presence-Absence PCoA

Jaccard distances between presence-absence of taxa were calculated using the *distance* function of the R package *philentropy* v0.2.0 [67]. The PCoA was generated using the *wcmdscale* function in *vegan* v2.5-3 [63]. The two best sampled genera, *Halorubrum* (orange) and *Haloferax* (red), are colored distinctively.

### Recognition Site Assignment

To determine the most likely recognition sites, each member of each cHG was searched against the REBASE Gold Standard set using BLASTp. The REBASE gold standard set was chosen over the individual gene sets on account of it having a much higher density of recognition site annotation. This simplifies the need to search for secondary hits to find predicted target sites. After applying an e-value cut-off of 1E-20, the top hit was assigned to each ORF.

**CTAG and GATC** motifs were counted with an inhouse perl script available at the Gogarten-lab’s GitHub [68].

### Gene Ontology

Sets of GO terms were identified for each cHG using Blast2GO [69]. Annotations were checked against the UniProt database [70] using arCOG identifiers.

## 3. Results

### RM-system gene distribution

Analysis of 217 haloarchaeal genomes and metagenome assembled genomes yielded 48 total candidate collapsed homologous groups (cHGs) of RM-system components. Out of these 48 cHGs, 26 had arCOG annotation suggesting DNA methylase activity, restriction enzyme activity, or specificity module activity as part of an RM system. We detected 22 weaker candidates with predominant arCOG annotations matching other functions (**Table 1**). Our analysis shows that nearly all of the cHGs are found more than once. (**Figure 1A**). Indeed, 16 families are found in 20 or more genomes each (>9 %), and this frequency steadily increases culminating in five families being conserved in greater than 80 genomes each (>37 %) with one cHG being in ∼80 % of all Haloarchaea surveyed. Though these genes appear frequently in taxa across the haloarchaeal class, the majority of each candidate RM system cHG is present in fewer than half the genomes, - the second most abundantly recovered cHG is found in only ∼47 % of all taxa surveyed. We note that the cHGs with wide distribution are annotated as MTases without an identifiable co-evolving restriction endonuclease: Group U DNA_methylase-022; W dam_methylase-031; Y dcm_methylase-044; and AT Uncharacterized-032 (members of this cHG are also annotated as methylation subunit and N6-Adenine MTase). Rarefaction analysis indicates about 50 % of the genomes assayed contain seven dominant cHGs, and that all taxa on average are represented by half of the cHGs (**Figure 1B**). Together, the separate analyses indicate extensive gene gain and loss of RM-system genes. In contrast, orphan MTases in cHG U and W, and to a lesser extent Y (Figure 2) have a wider distribution in some genera (see below for further discussion).

**Figure 1:**
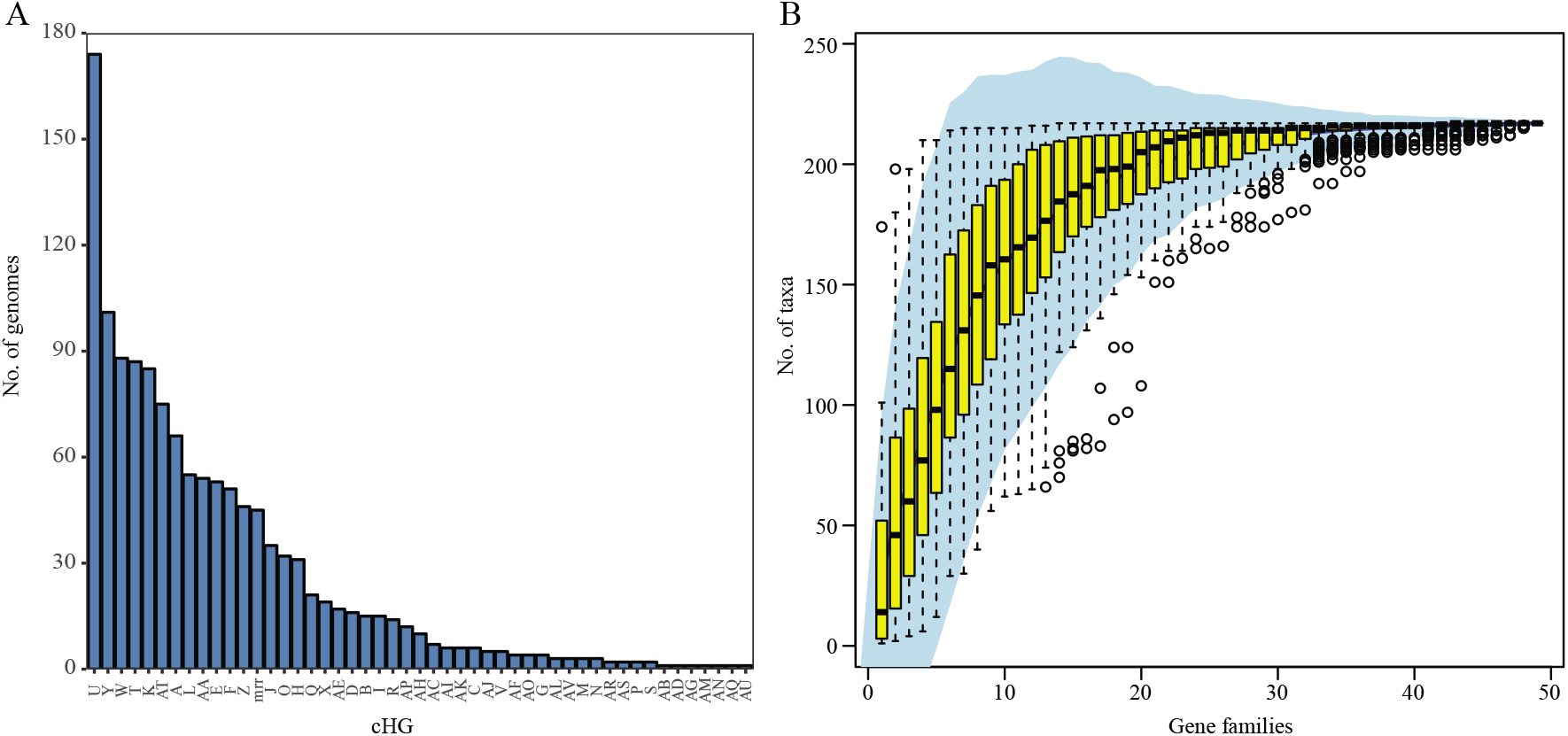
Distribution of collapsed Homologous Group (cHG) among haloarchaeal genomes. (A) the number of genomes present in each collapsed Homologous Group (cHG). No cHG contains a representative from every genome used in this study. With the exception of one cHG, all contain members from fewer than half of the genomes. The cHGs are ordered by number of genomes they contain. (B) rarefaction plot of the number of genomes represented as cHGs accumulate. 95% confidence interval is shown in shaded blue area and yellow box whisker plots give the number of taxa from random subsamples (permutations = 100) over 48 gene families.

**Figure 2:**
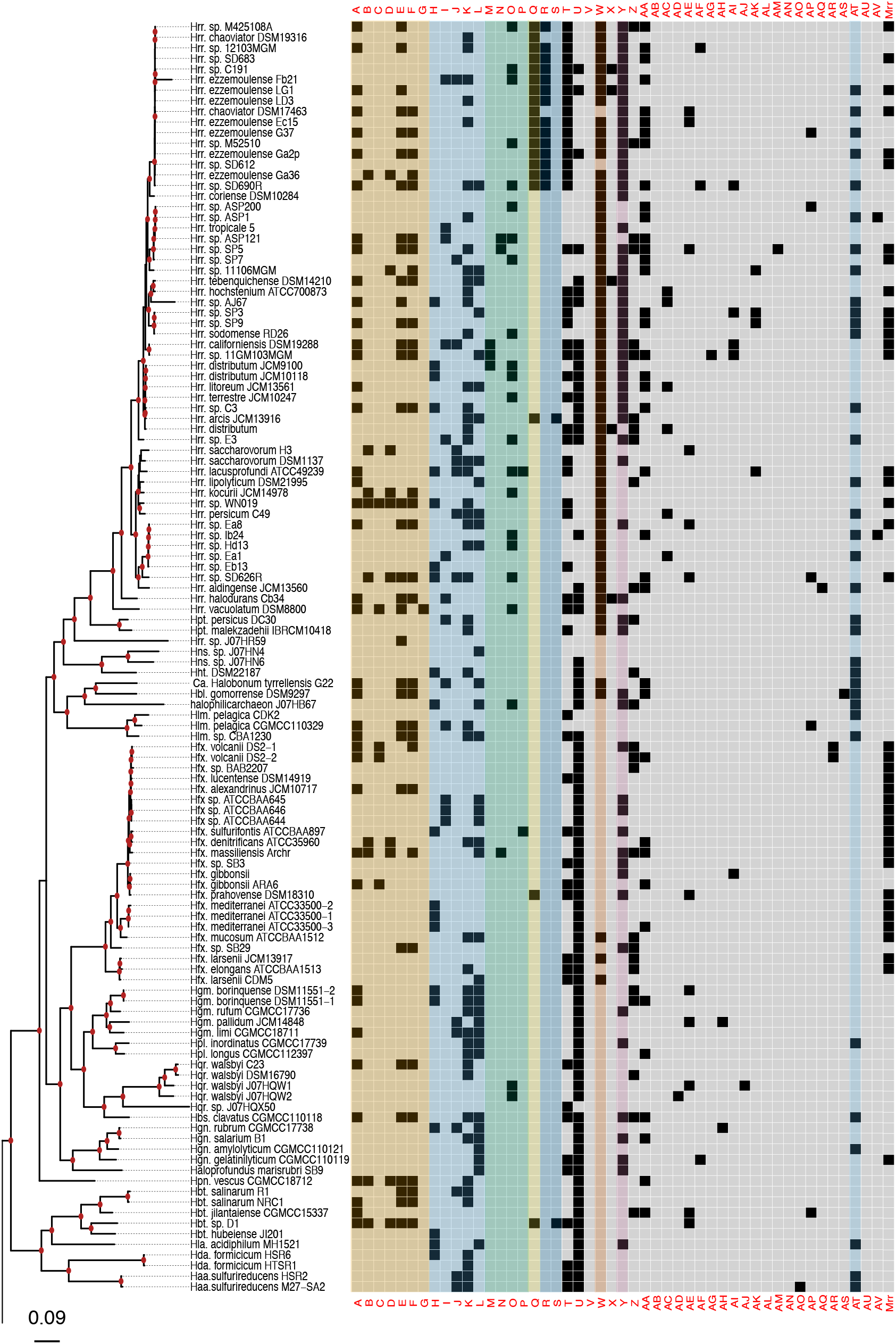
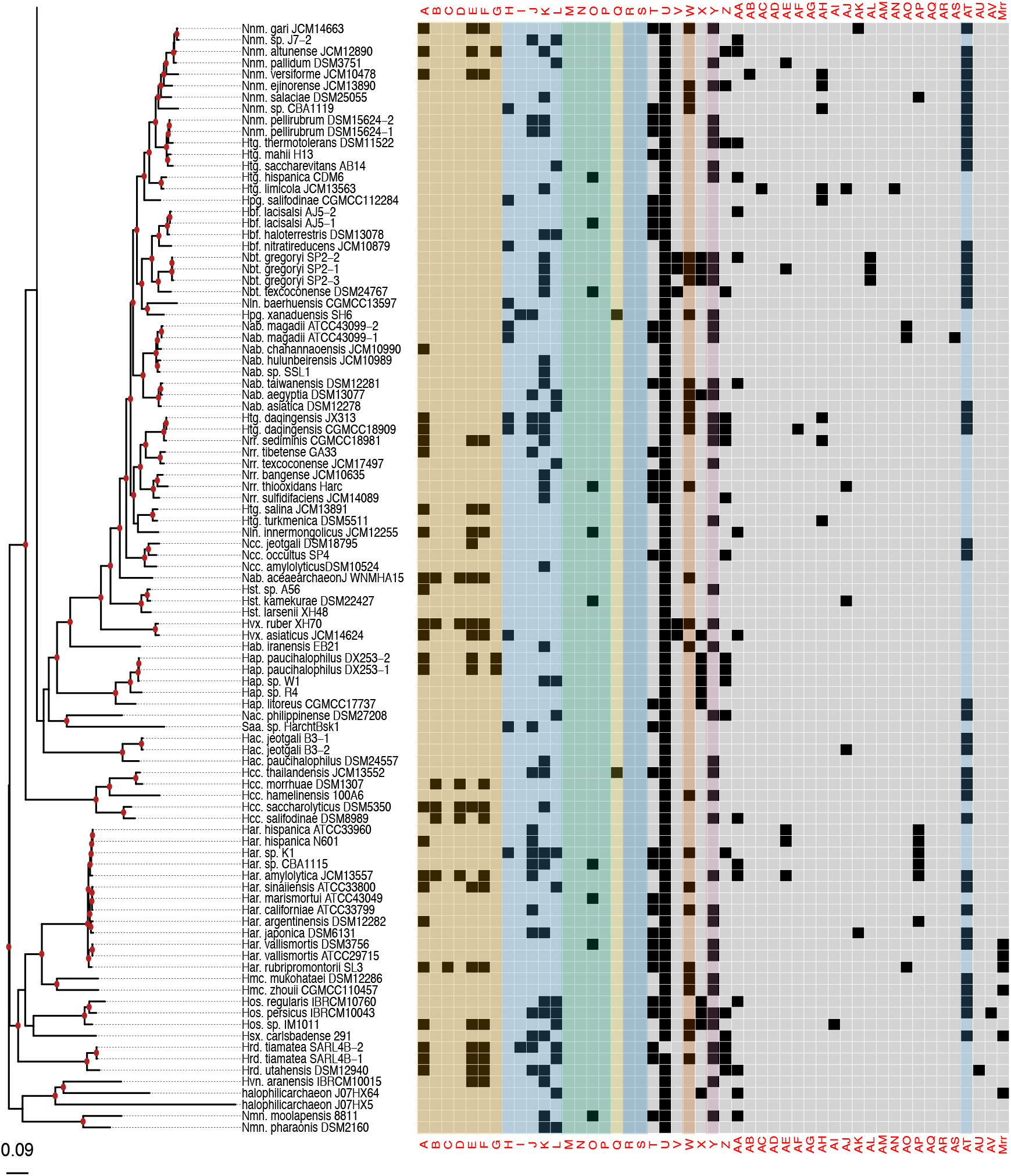
Presence-absence matrix of the 48 candidate RMS cHGs plotted against the reference phylogeny. For most cHGs the pattern of presence-absence does not match the reference phylogeny (compare supplementary Figures S2-S5) RMS-candidate cHGs are loosely ordered by system type and with the ambiguously assigned RM candidates at the end. Above is a key relating the column names to the majority functional annotation.

**Table 1.**
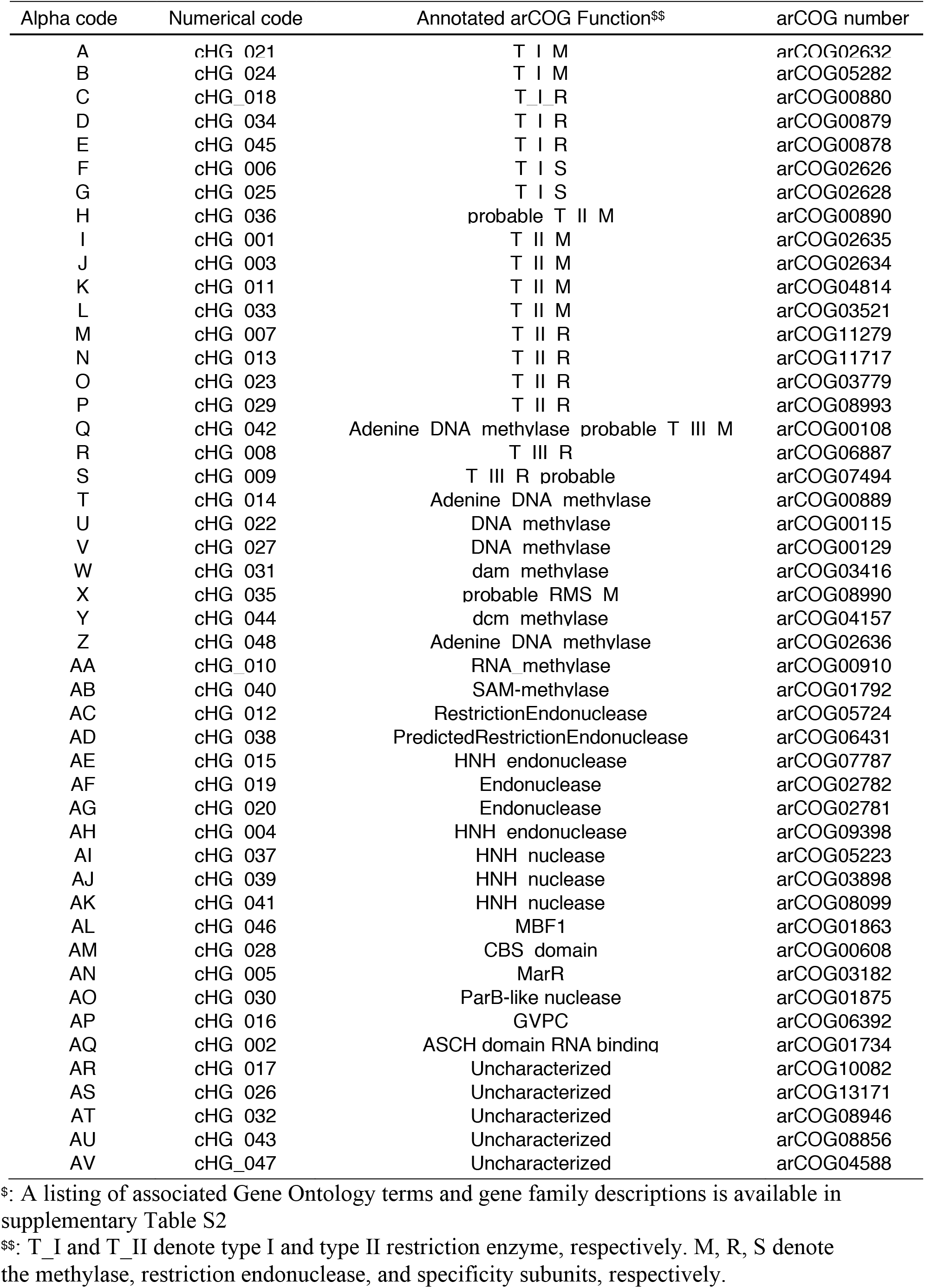
Collapsed homologous group descriptions.^$^

| Alpha code | Numerical code | Annotated arCOG Function\$\$ | arCOG number |
| --- | --- | --- | --- |
| A | cHG_021 | T_I_M | arCOG02632 |
| B | cHG_024 | T_I_M | arCOG05282 |
| C | cHG_018 | T_I_R | arCOG00880 |
| D | cHG_034 | T_I_R | arCOG00879 |
| E | cHG_045 | T_I_R | arCOG00878 |
| F | cHG_006 | T_I_S | arCOG02626 |
| G | cHG_025 | T_I_S | arCOG02628 |
| H | cHG_036 | probable_T_II_M | arCOG00890 |
| I | cHG_001 | T_II_M | arCOG02635 |
| J | cHG_003 | T_II_M | arCOG02634 |
| K | cHG_011 | T_II_M | arCOG04814 |
| L | cHG_033 | T_II_M | arCOG03521 |
| M | cHG_007 | T_II_R | arCOG11279 |
| N | cHG_013 | T_II_R | arCOG11717 |
| O | cHG_023 | T_II_R | arCOG03779 |
| P | cHG_029 | T_II_R | arCOG08993 |
| Q | cHG_042 | Adenine DNA methylase probable T_III_M | arCOG00108 |
| R | cHG_008 | T_III_R | arCOG06887 |
| S | cHG_009 | T_III_R probable | arCOG07494 |
| T | cHG_014 | Adenine DNA methylase | arCOG00889 |
| U | cHG_022 | DNA methylase | arCOG00115 |
| V | cHG_027 | DNA methylase | arCOG00129 |
| W | cHG_031 | dam methylase | arCOG03416 |
| X | cHG_035 | probable RMS_M | arCOG08990 |
| Y | cHG_044 | dcm methylase | arCOG04157 |
| Z | cHG_048 | Adenine DNA methylase | arCOG02636 |
| AA | cHG_010 | RNA methylase | arCOG00910 |
| AB | cHG_040 | SAM-methylase | arCOG01792 |
| AC | cHG_012 | RestrictionEndonuclease | arCOG05724 |
| AD | cHG_038 | PredictedRestrictionEndonuclease | arCOG06431 |
| AE | cHG_015 | HNH endonuclease | arCOG07787 |
| AF | cHG_019 | Endonuclease | arCOG02782 |
| AG | cHG_020 | Endonuclease | arCOG02781 |
| AH | cHG_004 | HNH endonuclease | arCOG09398 |
| AI | cHG_037 | HNH nuclease | arCOG05223 |
| AJ | cHG_039 | HNH nuclease | arCOG03898 |
| AK | cHG_041 | HNH nuclease | arCOG08099 |
| AL | cHG_046 | MBF1 | arCOG01863 |
| AM | cHG_028 | CBS domain | arCOG00608 |
| AN | cHG_005 | MarR | arCOG03182 |
| AO | cHG_030 | ParB-like nuclease | arCOG01875 |
| AP | cHG_016 | GVPC | arCOG06392 |
| AQ | cHG_002 | ASCH domain RNA binding | arCOG01734 |
| AR | cHG_017 | Uncharacterized | arCOG10082 |
| AS | cHG_026 | Uncharacterized | arCOG13171 |
| AT | cHG_032 | Uncharacterized | arCOG08946 |
| AU | cHG_043 | Uncharacterized | arCOG08856 |
| AV | cHG_047 | Uncharacterized | arCOG04588 |

\$: A listing of associated Gene Ontology terms and gene family descriptions is available in supplementary Table S2
\$\$: T\_I and T\_II denote type I and type II restriction enzyme, respectively. M, R, S denote the methylase, restriction endonuclease, and specificity subunits, respectively.

The phylogeny of the class Halobacteria inferred from concatenated ribosomal proteins (**Figure 2**) was largely comparable to prior work [71], and with a taxonomy based on concatenations of conserved proteins [72,73]. For instance, in our phylogeny the *Halorubracaea* group with the *Haloferacaceae* recapitulating the order *Haloferacales*, and the families, *Halobacteriaceae, Haloarculaceae* and *Halococcaceae* group within the order *Halobacteriales*. Our genome survey in search of RM-system genes encompassed a broad taxonomic sampling, and it explores in depth the genus *Halorubrum* because it is a highly speciated genus, and because the existence of many genomes from the same species allows within species distribution assessment.

Comparison of the phylogeny in **Figure 2** to the heatmap giving the presence/absence of RM system cHG candidates demonstrates that the cHG distribution is highly variable (**Figure 2**). The one glaring exception is cHG U, a DNA methylase found in 174 of the 217 genomes analyzed. Since it is not coupled with a restriction enzyme of equal abundance, it is assumed to be an orphan MTase. The MTase from *Hfx. volcanii* (gene HVO_0794), which recognizes the CTAG motif [45] is a member of this cHG. Though U is widely distributed, within the genus *Halorubrum* it is only found in ∼37.5 % (21/56) of the genomes. While U’s phylogenetic profile is compatible with vertical inheritance over much of the phylogeny, the presence absence data also indicate a few gene transfer and loss events within *Halorubrum*. cHG U is present in *Hrr. tebenquichense* DSM14210, *Hrr. hochstenium* ATCC700873, *Hrr*. sp. AJ767, and in strains from the related species *Hrr. distributum, Hrr. arcis, Hrr. litoreum* and *Hrr. terrestre* suggesting an acquisition in the ancestor of this group.

Instead of U, another orphan MTase is abundantly present in *Halorubrum* spp., cHG W. It was found in ∼95 % of all *Halorubrum* strains, with three exceptions - an assembled genome from metagenome sequence data, and two from incomplete draft genomes of the species *Halorubrum ezzemoulense*. Interestingly, when U is present in a *Halorubrum* sp. genome, so too is W (**Figure 2**). In a complementary fashion, analysis of W outside of the *Halorubrum* shows that it is found patchily distributed throughout the rest of the class Halobacteria (∼20 % -32/158), and always as a second orphan MTase with cHG U. When the members of cHG W were used to search the uniprot database, the significant matches included the *E. coli* Dam MTase, a very well-characterized GATC MTase, which provides strong evidence that this cHG is a GATC orphan MTase family. The presence and absence of cHG U and W in completely sequenced genomes is given in Table S3, together with the frequency of the CTAG and GATC motifs in the main chromosome.

The rest of the RM cHGs are much more patchily distributed (**Figure 2**). For instance, the cHGs that make up columns A-G represent different gene families within the Type I RM system classification; two MTases (A,B), three REases (C,D,E), and two site specificity units (SSUs) (F,G). Throughout the Haloarchaea, cHGs from columns A, E and F, representing an MTase, an REase, and an SSU respectively, are found co-occurring 35 times. In a subset of genomes studied for synteny, A, E, and F are encoded next to one another in *Natrinema gari, Halorhabdus utahensis, Halorubrum* SD690R, *Halorubrum ezzemoulense* G37, and *Haloorientalis* IM1011 (**Figure 3**). These genes probably represent a single transcriptional unit of genes working together for restriction and modification purposes. Since the Type I RM system is a five-component system, the likely stoichiometry is 2:2:1. These three cHGs co-occur four times within the species *Halorubrum ezzemoulense*, and two of these cHGs (A and E) co-occur an additional three more times, suggesting either a loss of the SSU, or an incomplete genome sequence for those strains. If it is due to incomplete sequencing, then 7/16 (43 %) of the *Hrr. ezzemoulense* genomes have this set of co-occurring genes, while half do not have an identified Type I system. This is particularly stunning since strains FB21, Ec15, G37 and Ga2p were all cultivated at the same time from the same sample, a hypersaline lake in Iran. Furthermore, one strain, Ga36, has a different identified Type I RM system composed of substituted cHGs A and E with B and D, respectively, while maintaining the same SSU. This suggests the same DNA motif may be recognized by the different cHGs and that these cHGs are therefore functionally interchangeable. Members of cHGs B, F, and D were found as likely co-transcribed units in *Halococcus salifodinae, Natronolimnobius aegyptiacus, Halorubrum kocurii, Haloarcula amylolytica* (**Figure 3**). In *Halorubrum* DL, and *Halovivax ruber* XH70, genomes that contained members from cHGs A, B, D, E, and F these genes were not found in a single unit, suggesting that they do not form a single RM system. Together, these analyses suggest this Type I RM system has a wide but sporadic distribution, that this RM system is not required for individual survival, and that functional substitutions occur for cHGs.

**Figure 3.**
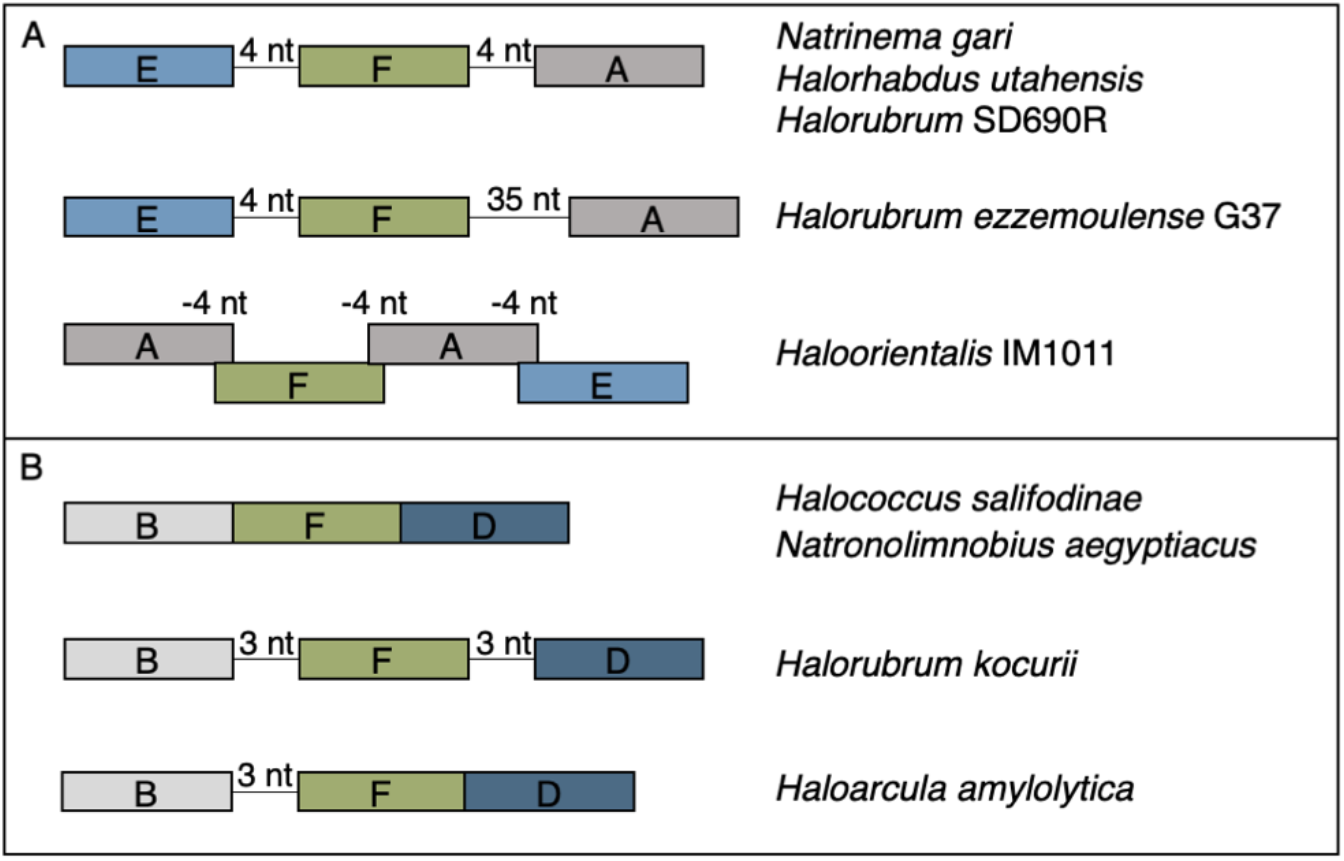
Gene maps for syntenic clusters of gene families (A) EFA and (B) BFD found in a subset of organisms identified to the right of each map. Genes are colored by gene families with Type I methylases (AB) in greys, Type I restriction endonucleases (DE) in blues, and Type I site specificity unit (F) in green.

Type II RM systems contain an MTase and an REase that target the same motif but do not require an associated SSU because each enzyme has its own TRD. The Type II RM system cHGs are in columns H-L for the MTases, and M-P for the REases. Memberships to the Type II MTase cHGs are far more numerous in the Haloarchaea than their REase counterpart, as might be expected when witnessing decaying RM systems through the loss of the REase. The opposite result, more REases is a more difficult scenario because an unmethylated host genome would be subject to restriction by the remaining cognate REase (e.g., addiction cassettes). There are 14 “orphan” Type II REases in **Figure 2**, but their cognate MTase’s absence could be explained by incomplete genome sequence data.

Type III RM systems have been identified in cHGs Q (MTase) and R and S (REases). Type III MTases and REases (cHGs Q and R) co-occur almost exclusively in the species *Halorubrum ezzemoulense*, our most highly represented taxon. Furthermore, these Type III RM systems are highly restricted in their distribution to that species, with cHGs co-occuring only twice more throughout the Haloarchaea, and with a different REase cHG (S); once in *Halorubrum arcis*, and another in *Halobacterium* D1. Orphan MTases occurred twice in cHG Q. Of particular interest is that closely related strains also cultivated from Lake Bidgol in Iran but which are in a different but closely related *Halorubrum* species (e.g., Ea8, IB24, Hd13, Ea1, Eb13) do not have a Type III RM system, implying though exposed to the same halophilic viruses, they do not rely on this system for avoiding virus infection.

Mrr is a Type IV REase that was suggested to cleave methylated GATC sites [74,75]. Mrr homologs are identified in most *Haloferax* sp., they have a sporadic distribution among other *Haloferacaceae* and in <u>the *Halobacteriaceae*</u>, and are absent in the *Natrialbaceae* (**Figure 2**). cHGs Z-AV are not sufficiently characterized to pinpoint of their role in DNA RM systems or as MTase. These cHGs likely include homing endonucleases or enzymes modifying nucleotides in RNA molecules; however, their function as orphan MTases or restriction endonucleases can, at present, not be excluded.

### Horizontal Gene Transfer explains patchy distribution

The patchy appearance of RM system candidates was further investigated by plotting the Jaccard distance of the presence-absence data against the alignment distance of the reference tree (supplementary **Figure S2**). If the presence-absence data followed vertical descent one would expect the best-fit line to move from the origin with a strong positive slope. Instead, the best fit line is close to horizontal with an r-squared value of 0.0047, indicating negligible relationship between the overall genome phylogeny and RM system complement per genome. The presence-absence clustering patterns were visualized by plotting a principle coordinate analysis (supplementary **Figure S3**). The high degree of overlap between the ranges of the three groups illustrates that there are few RM system genes unique to a given group and a large amount of overlap in repertoires.

To further evaluate the lack of long-term vertical descent for RM system genes, a phylogeny was inferred from the presence-absence pattern of cHGs. The resultant tree (**Figure S4**) is largely in disagreement with the reference phylogeny. The bootstrap support set from the presence-absence phylogeny was mapped onto the ribosomal topology (**Figure S5**). The resulting support values demonstrate an extremely small degree of agreement between the two methods. The few areas where there is even 10% support are near the tips of the ribosomal phylogeny and correspond to parts of established groups, such as *Haloferax, Natronobacterium*, and *Halorubrum*. Internode Certainty (IC) scores are another way to compare phylogenies. An average IC score of 1 represents complete agreement between the two phylogneies, and score of −1 complete disagreement. The average IC scores for the reference tree using the support set from the F81 tree was −0.509, illustrating that the presence absence data do not support the topology of the reference phylogeny.

The patchy distribution of the RM system candidate genes and their lack of conformity to the reference phylogeny suggests frequent horizontal gene transfer combined with gene loss events as the most probable explanation for the observed data. To quantify the amount of transfer, the TreeFix-Ranger pipeline was employed. TreeFix-DTL resolves poorly supported areas of gene trees to better match the concatenated ribosomal protein gene tree used as reference. Ranger-DTL resolves optimal gene tree rooting against the species tree and then computes a reconciliation estimating the number of duplications, transfers, and losses that best explains the data (**Table 2**). For almost every cHG with four or more taxa, our analysis infers several HGT events. Only cHG R, a putative Type III restriction enzyme found only in a group of closely related *Halorubrum ezzemoulense strains*, has not been inferred to undergo at least one transfer event.

**Table 2.**
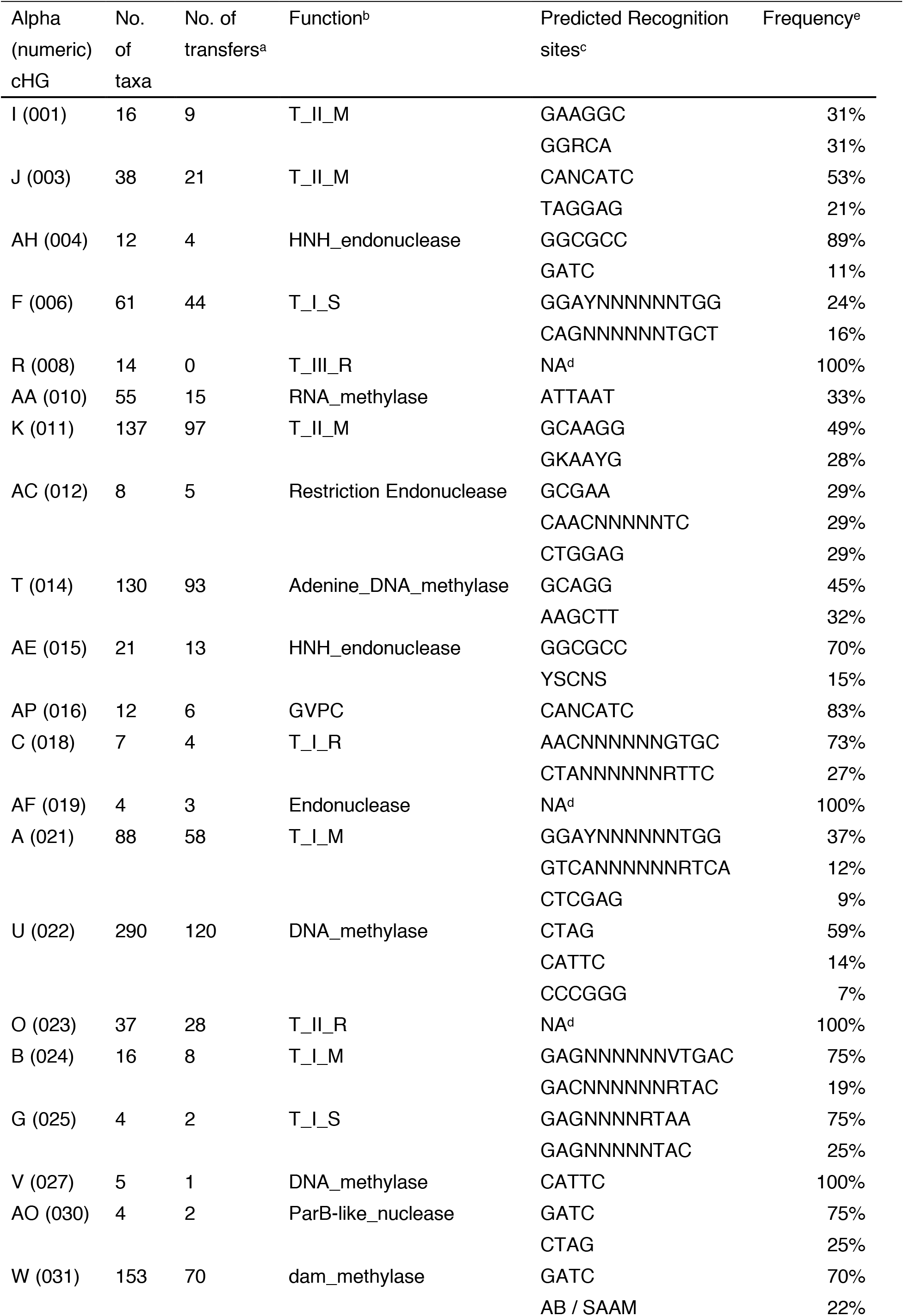
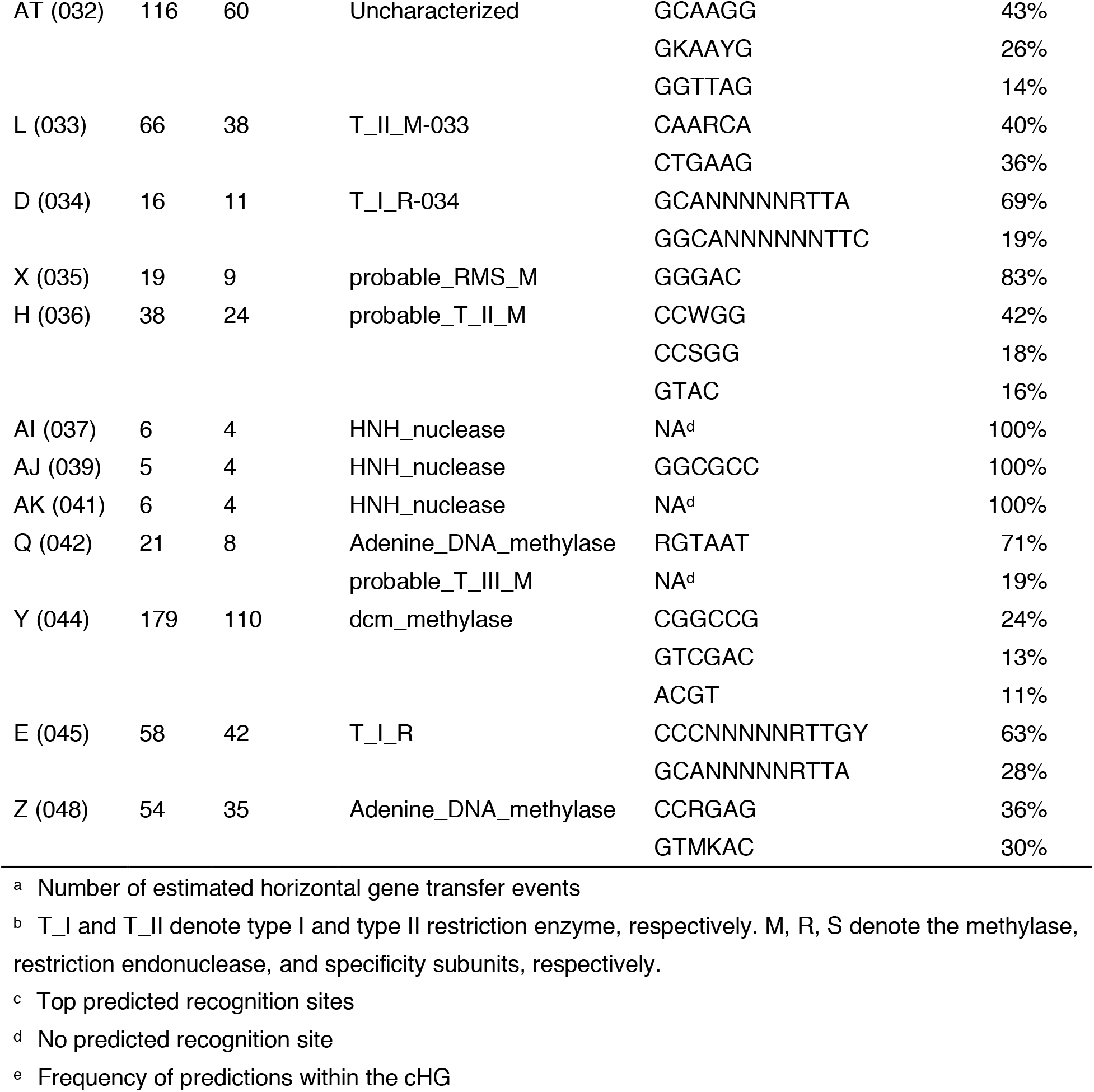
Important traits of cHGs with four or more ORFs.

RM systems usually function as cooperative units [48,76,77]. It stands to reason that some of the RM system candidates may be transferred as units, maintaining their cognate functionality. This possibility was examined by a correlation analysis. A spearman correlation was made between all pairs of cHGs. Those with a significant result at a Bonferroni-corrected p <0.05 were plotted in a correlogram (**Figure 4**). As illustrated in **Figure 3**, cHGs with significant similar phylogenetic profiles often are near to one another in the genomes.

**Figure 4.**
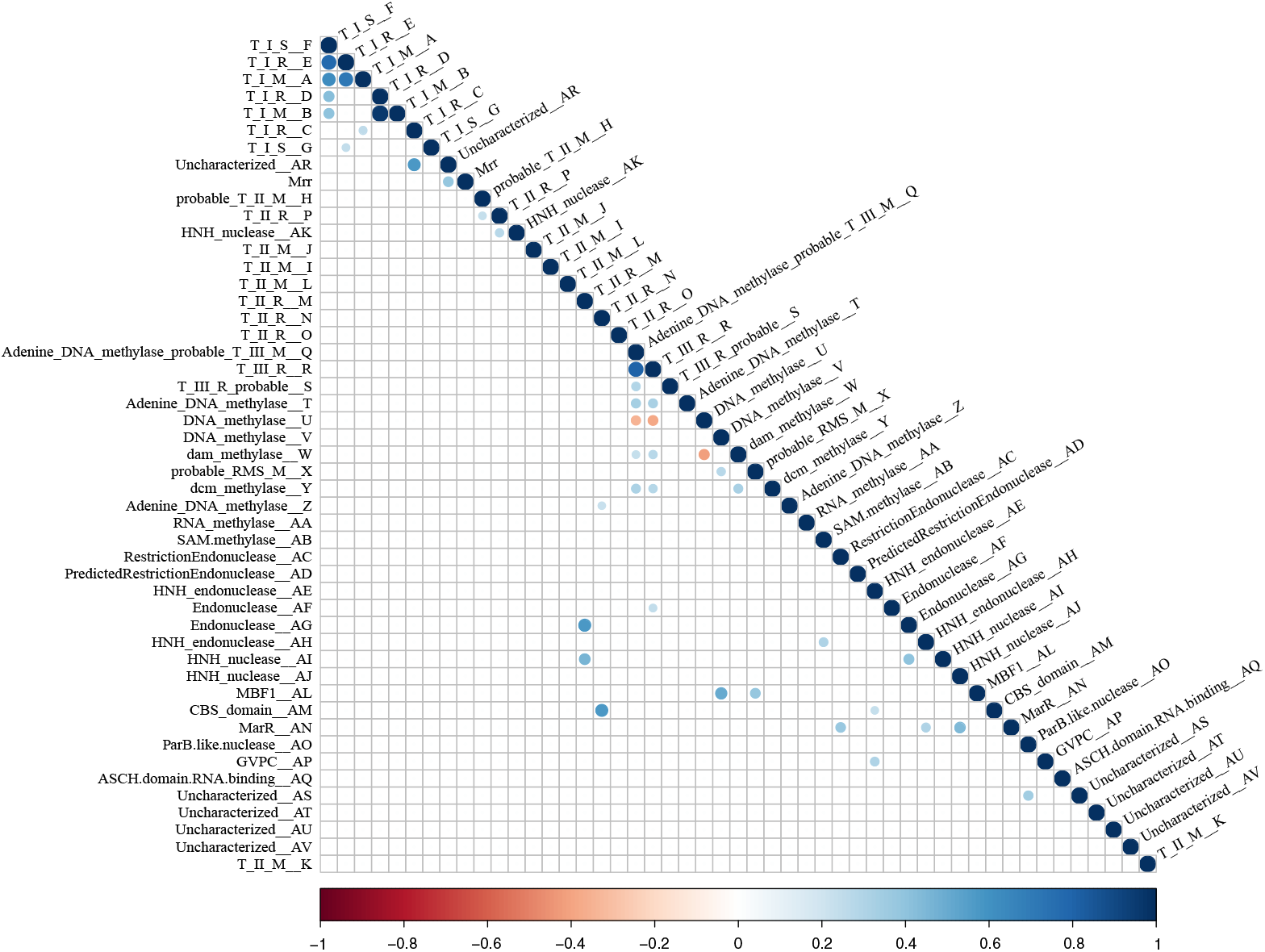
Heatmap of co-occurrence between the 48 RMS-candidate cHGs. Positive correlation indicates the cHGs co-occur while negative indicates that the presence of one means the other will not be present. Significance level is p < 0.05 with a Bonferroni correction applied for multiple tests. Blue indicates significant positive correlation; red indicates a significant negative correlation.

## 4. Discussion

A striking result of our study is the irregular distribution of the RM system gene candidates throughout not just the haloarchaeal class, but also within its orders, genera, species, and even communities and populations. The patchy distribution is almost certainly the result of frequent HGT and gene loss. RM system genes are well known for their susceptibility to HGT and loss, and their presence almost never define a clade or an environmental source (e.g., [36,78]). Frequent acquisition of RM system genes through HGT is illustrated by their sporadic distribution. For example, *Halorubrum* genomes encode many candidate RM system cHGs that are absent from the remainder of the Halobacteria (e.g., cHG M, R, S, AC, AG, AM). Only one of these (cHG R) is found in more than 3 genomes, a Type III restriction protein found in 14 of 57 *Halorubrum* genomes. Mrr homologs have a sporadic distribution among *Haloferacaceae* and <u>*Halobacteriaceae*</u> and are absent in *Natrialbaceae* (**Figure 2**). Gene loss undoubtedly contributed to the sparse cHGs distribution; however, without invoking frequent gene transfer, many independent and parallel gene losses need to be postulated. We also observed that a number haloarchaeal species possess multiple Type I subunit genes, allowing for functional substitution of the different subunits in the RM system. The existence of multiple Type I subunits has also been observed in *Helicobacter pylori*, in which 4 different SSU loci are used by the organism’s Type I system to target different recognition sequences; these SSUs can even exchange TRDs, resulting in variation in the methylome of *H. pylori* [79–81]. In our results, however, we observed multiple MTase and REase subunits alongside a single SSU, suggesting the functional substitution of the subunits in these haloarchaeal organisms does not result in variation in detected recognition sequences.

Mrr is a Type IV REase that cleaves methylated target sites. Studies have demonstrated that this gene reduces transformation efficiency of GATC-methylated plasmids in *H. volcanii*, and that deletion of the *mrr* gene increases transformation efficiency on GATC-methylated plasmids, suggesting that this Type IV REase can target GATC-methylated sites for cleavage [74,75]. However, we find no anticorrelation between the presence of Mrr homologs and members of cHG W, which is homologous to the *E. coli* Dam MTase, a very well-characterized GATC MTase (**Figures 2 and 4**). This suggests that some members of cHG W or the Mrr homologs either are dysfunctional of have a site specificity different from the GATC motif.

It seems counterintuitive that RM systems are not more conserved as cellular countermeasures against commonly occurring viruses. It may be that cells do not require extensive protection via RM systems, because they use multiple defensive systems some of which might be more effective. For example, another well-known defense against viruses is the CRISPR-Cas system [82]. CRISPR recognizes short (∼40bp) regions of invading DNA that the host has been exposed to previously and degrades it. While it can be very useful against virus infection, our prior work indicated that CRISPR-Cas was also sporadically distributed within communities of closely related haloarchaeal species [83] indicating they are not required for surviving virus infection.

Both the RM and CRISPR-Cas systems are only important countermeasures after external fortifications have failed to prevent a virus from infiltrating, and therefore their limited distributions also indicate that the cell’s primary defense would be in preventing virus infection altogether, which is accomplished by different mechanisms. By altering surfaces via glycosylation cells can avoid virus predation prior to infection. In *Haloferax* species there are two pathways which control glycosylation of external features. One is relatively conserved and could have functions other than virus avoidance, while the other is highly variable and shows hallmarks of having genes mobilized by horizontal transfer [84]. At least one halovirus has been found to require glycosylation by its host in order to infect properly [85]. Comparison of genomes and metagenomes from hypersaline environments showed widespread evidence for distinct “genomic” islands in closely related halophiles [86] that contain a unique mixture of LPS and other genes that contribute to altering the cell’s surface structure and virus docking opportunities. Thus selective pressure on post infection, cytosolic and nucleic acids-based virus defenses is eased, allowing them to be lost randomly in populations.

A major consideration in understanding RM system diversity is that viruses, or other infiltrating selfish genetic elements, might gain access to the host’s methylation after a successful infection that was not stopped by the restriction system. Indeed, haloviruses are known to encode DNA methyltransferases in their genomes (e.g., see [87]). In this case, RM systems having a limited within population distribution would then be an effective defense for that part of the population possessing a different RM system. Under this scenario, a large and diverse pool of mobilized RM system genes could offer a stronger defense for the population as a whole. A single successful infection would no longer endanger the entire group of potential hosts.

Group selection may be invoked to explain the within population diversity of RM systems; a sparse distribution of RM systems may provide a potential benefit to the population as a whole, because a virus cannot easily infect all members of the population. However, often gene level selection is a more appropriate alternative to group selection [88,89]. Under a gene centered explanation, RM systems are considered as selfish addiction cassettes that may be of little benefit to its carrier. While RM systems may be difficult to delete as a whole, stepwise deletion, that begins with inactivation of the REase activity can lead to their loss from a lineage. Their long-term survival thus may be a balance of gain through gene transfer, persistence through addiction, and gene loss. This gene centered explanation is supported by a study from [36], which examined the distribution of MTase genes in ∼1000 bacterial genomes. They observed, similar to our results in the Halobacteria, that MTases associated with RM systems are poorly conserved, whereas orphan MTases share conservation patterns similar to average genes. They also demonstrated that many RM-associated and orphan MTases are horizontally acquired, and that a number of orphan MTases in bacterial genomes neighbor degraded REase genes, suggesting that they are the product of degraded RM systems that have lost functional REases [36]. Similarly, Kong et al. [78] studying genome content variation in *Neisseria meningitidis* found an irregular distribution of RM systems, suggesting that these systems do not form an effective barrier to homologous recombination within the species. Kong et al. also observed that the RM systems themselves had been frequently transferred within the species. We conclude that RM genes in bacteria as well as archaea appear to undergo significant horizontal transfer and are not well-conserved. Only when these genes pick up additional functions, do parts of these systems persist for longer periods of time, as exemplified in the distribution of orphan MTases. However, the transition from RM system MTase to orphan MTase is an infrequent event. A study of 43 pan-genomes by Oliveira et al. [90] suggests that orphan MTases occur more frequently from transfer via large mobile genetic elements (MGEs) such as plasmids and phages rather than arise *de novo* from RM degradation. The distribution of orphan methylase cHG U and W, and their likely target motifs, CTAG and GATC, respectively suggests different biological functions for these two methylases. The widespread conservation of the CTAG MTase family cHG U supports the findings of Blow *et al*. [37] who identified a well-conserved CTAG orphan MTase family in the Halobacteria. Similar to other bacterial and archaeal genomes [91], the CTAG motif, the likely target for methylases in cHG U, is underrepresented in all haloarchaeal genomes (see table S3). The low frequency of occurrence, only about once per 4000 nucleotides, suggests that this motif and the cognate orphan methylase are not significantly involved in facilitating mismatch repair. The underrepresented CTAG motif was found to be less underrepresented near rRNA genes [91] and on plasmids; the CTAG motif also is a known target sequence for some IS elements [92]; and it may be involved in repressor binding, where the CTAG motif was found to be associated with kinks in the DNA when bound to the repressor [93,94] Interestingly, CTAG and GATC motifs are absent, or underrepresented in several haloarchaeal viruses [87,95,96]. Both, the presence of the cHG U methylase and the underrepresentation of the CTAG motif, appear to be maintained by selection; however, at present the reasons for the underrepresentation of the motif in chromosomal DNA, and the role that the methylation of this motif may play remain open questions.

## 5. Conclusions

RM systems have a sporadic distribution in haloarchaea, even within species and populations. In contrast, orphan methylases are more persistent in lineages, and the targeted motifs are under selection for lower (in case of CTAG) or higher (in case of GATC) than expected frequency. In case of the GATC motif, the cognate orphan MTase was found only in genomes where this motif occurs with high frequency.

## Supplementary Materials

The following are available online at www.mdpi.com/xxx/s1,

**Figure S1**. Workflow of RMS-candidate gene search strategy.

**Figure S2**. Plot of alignment distance as a function of presence-absence distance.

**Figure S3**. PCoA plot of the distances between the RMS presence-absence profiles of the 217 analyzed Halobacterial genomes.

**Figure S4**. Maximum-likelihood phylogeny of cHG presence-absence matrix.

**Figure S5**. Bootstrap support values of the presence-absence phylogeny mapped onto the ribosomal protein reference tree.

**Table S1**. Basic statistics for Halobacteriaceae complete and draft genomes

**Table S2**. Gene Ontology (GO) terms for each collapsed homologous group

**Table S3**. Distribution of orphan methylases cHGs U and W and frequency of their putative recognition motifs in completely sequenced halobacterial chromosomes.

## Author Contributions

Conceptualization, Thane R. Papke and Johann Peter Gogarten; Data curation, Matthew S. Fullmer, Matthew Ouellette and Artemis S. Louyakis; Formal analysis, Matthew S. Fullmer, Matthew Ouellette and Artemis S. Louyakis; Funding acquisition, Thane R. Papke and Johann Peter Gogarten; Investigation, Matthew S. Fullmer and Matthew Ouellette; Methodology, Matthew S. Fullmer, Artemis S. Louyakis and Johann Peter Gogarten; Project administration, Thane R. Papke and Johann Peter Gogarten; Software, Matthew S. Fullmer and Artemis S. Louyakis; Supervision, Thane R. Papke and Johann Peter Gogarten; Validation, Matthew S. Fullmer; Visualization, Matthew S. Fullmer and Artemis S. Louyakis; Writing – original draft, Matthew S. Fullmer and Johann Peter Gogarten; Writing – review & editing, Matthew Ouellette, Artemis S. Louyakis, Thane R. Papke and Johann Peter Gogarten.

## Acknowledgments

The Computational Biology Core, Institute for Systems Genomics, University of Connecticut provided computational resources. The authors thank an anonymous reviewer for pointing out the presence of Mrr in *Haloferax volcanii*.

