## Supplementary Tables and Figures for "The Patchy Distribution of Restriction-Modification System Genes and the Conservation of Orphan Methyltransferases in Halobacteria": Suppl_Figures.pdf (from another version of this preprint: version 1, 2019-02-15)

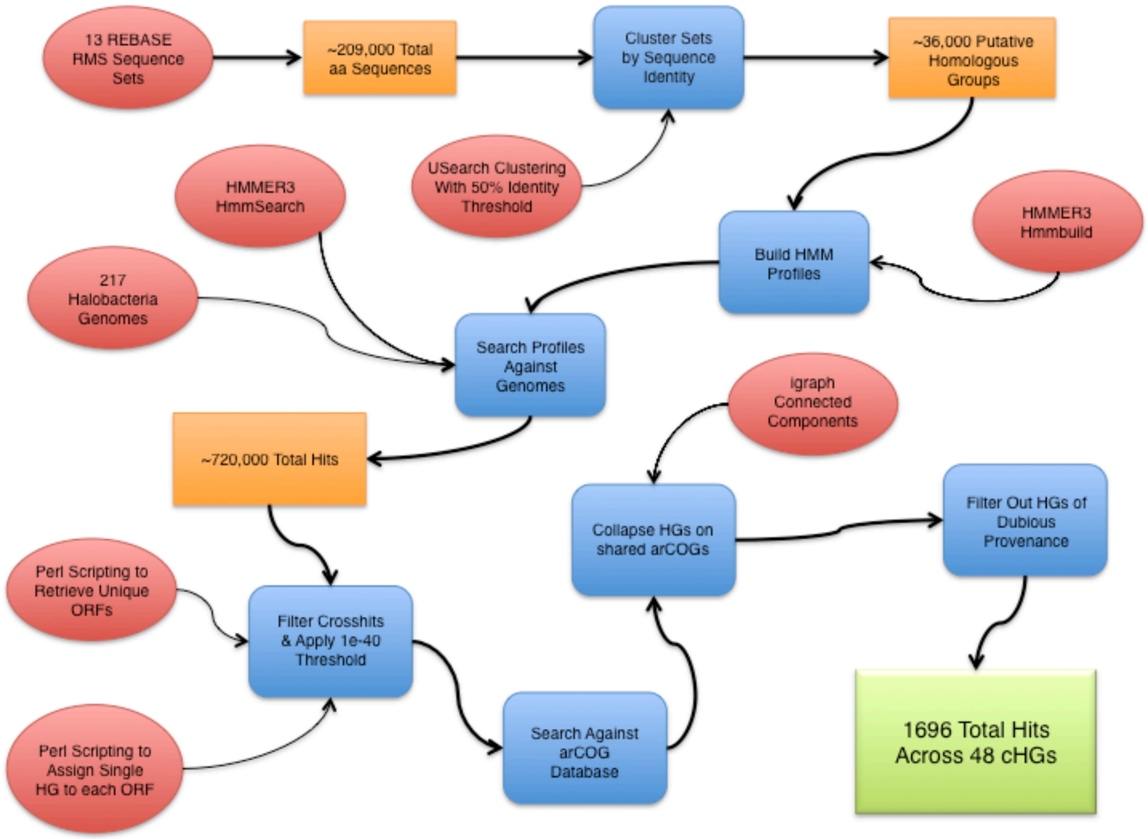


**Figure S1.** Workflow of RMS-candidate gene search strategy. Red circles represent input data and tools used to operate on data. Blue rectangles represent use and modification of the data during the process. Orange rectangles represent outputs at significant intermediate points in the process.

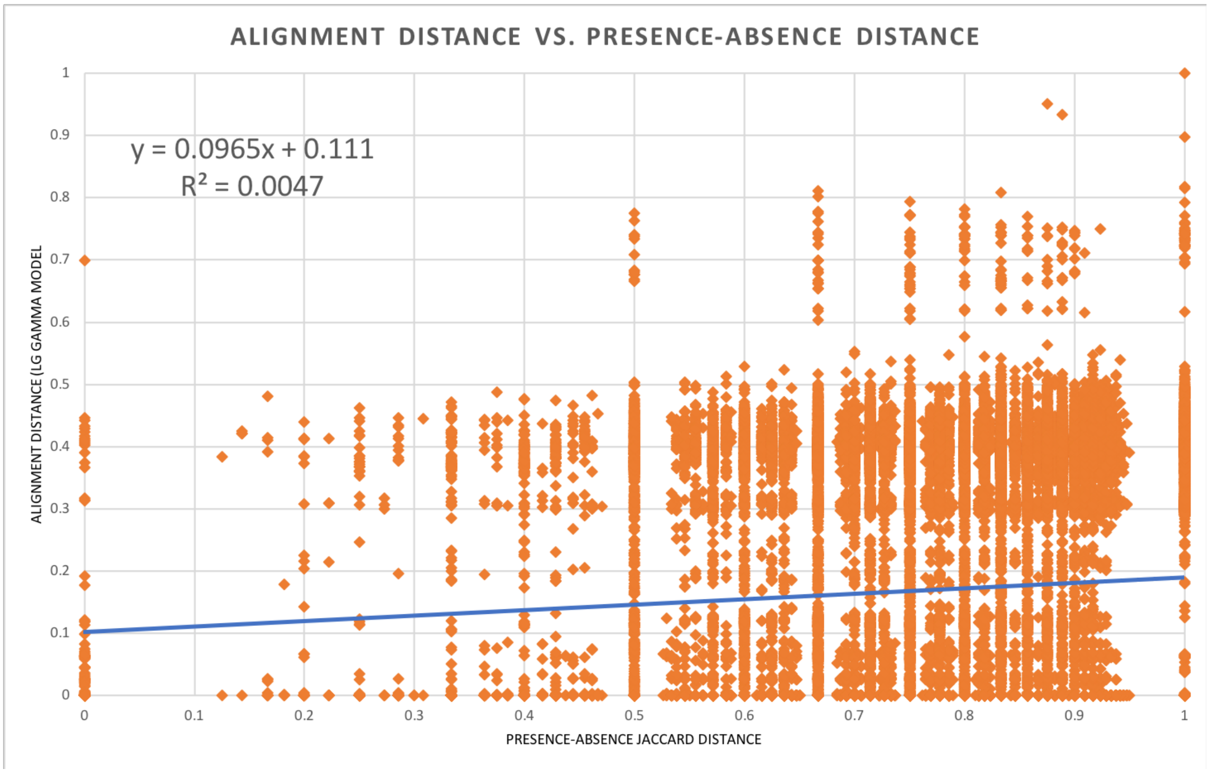


**Figure S2.** Plot of alignment distance as a function of presence-absence distance. Alignment distance was calculated from an LG+Gamma model and presence-absence using the Jaccard distance. If presence-absence pattern were a function of vertical descent, then the best fit line should broadly follow a line through the origin (0/0). The correlation between the two measures is negligible (r-squared = 0.0047).

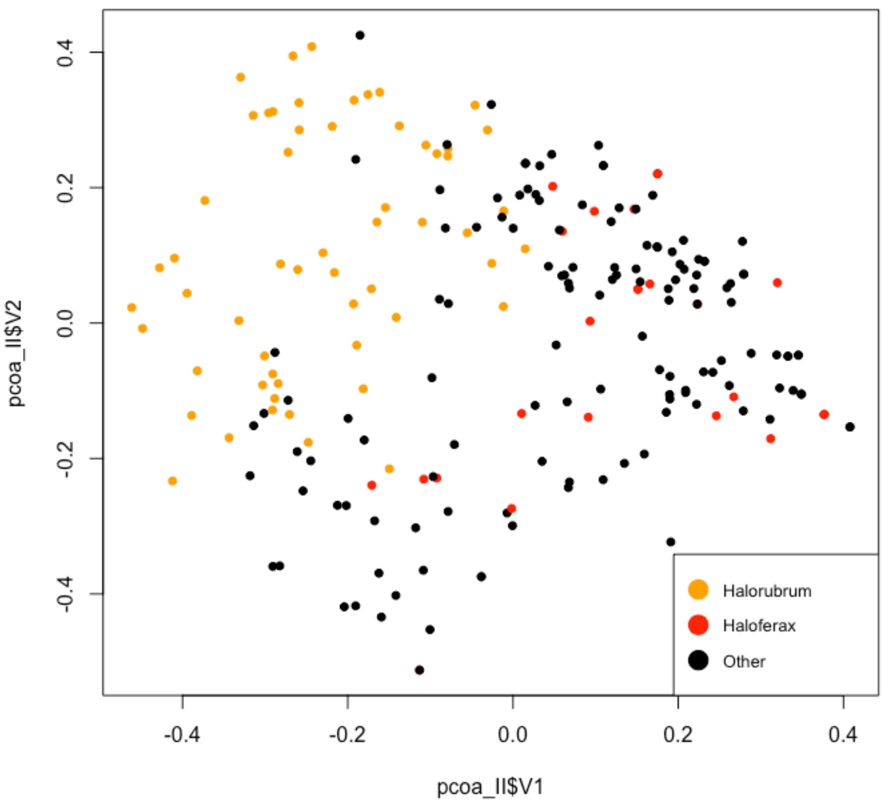


**Figure S3.** PCoA plot of the distances between the RMS presence-absence profiles of the 217 analyzed Halobacterial genomes. The two best sampled groups, *Halorubrum* (orange) and *Haloferax* (red), are colored separately. The overlap of ranges between the three groups illustrates that few RM system genes are unique to a given group and a large overlap in inventories between groups.

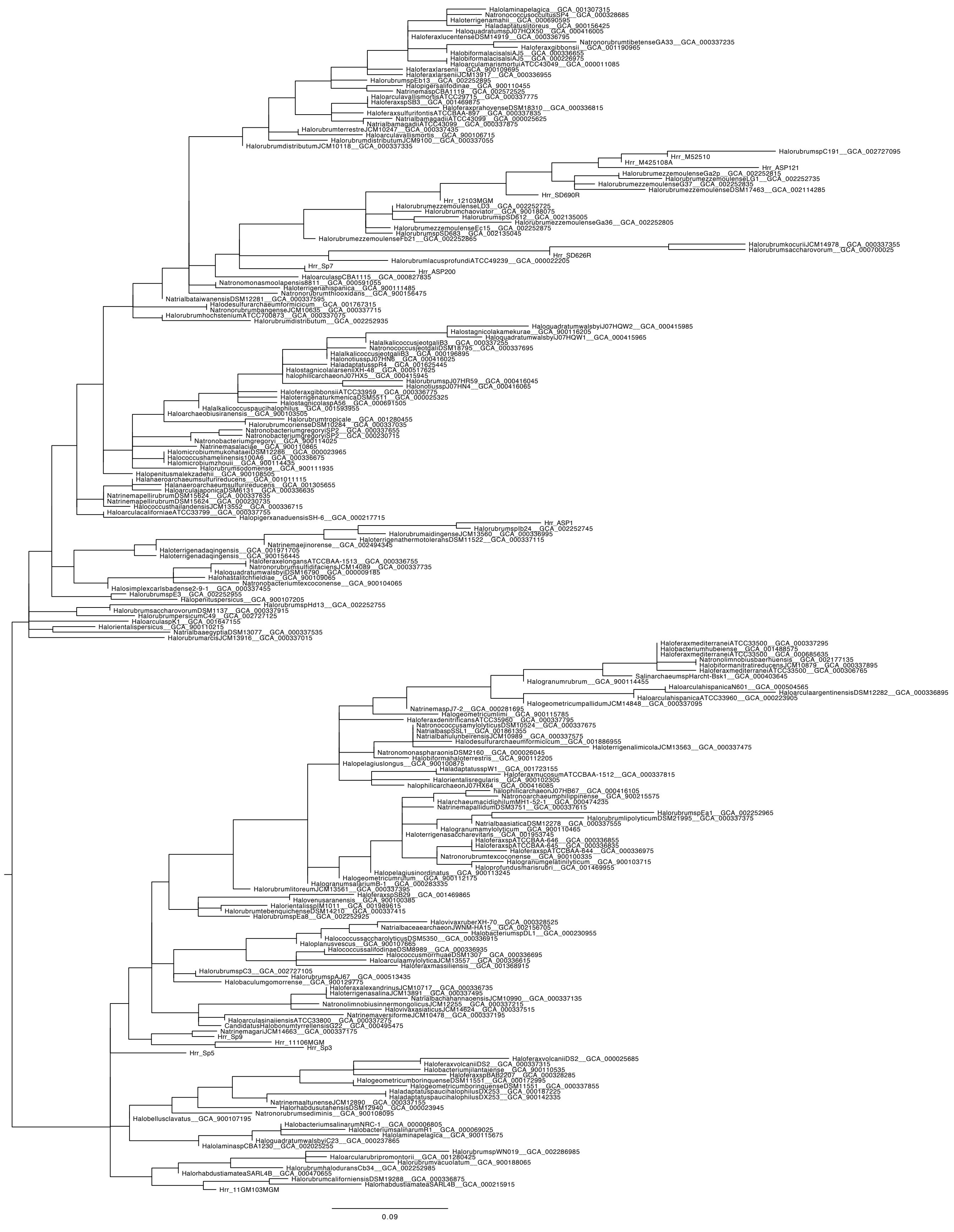
0.09

**Figure S4.** Maximum-likelihood phylogeny of cHG presence-absence matrix. An F81 model with empirical base-frequency was employed. The root was placed arbitrarily at the midpoint on account of the topology not bearing meaningful resemblance to the reference topology (cf. Figure 2).

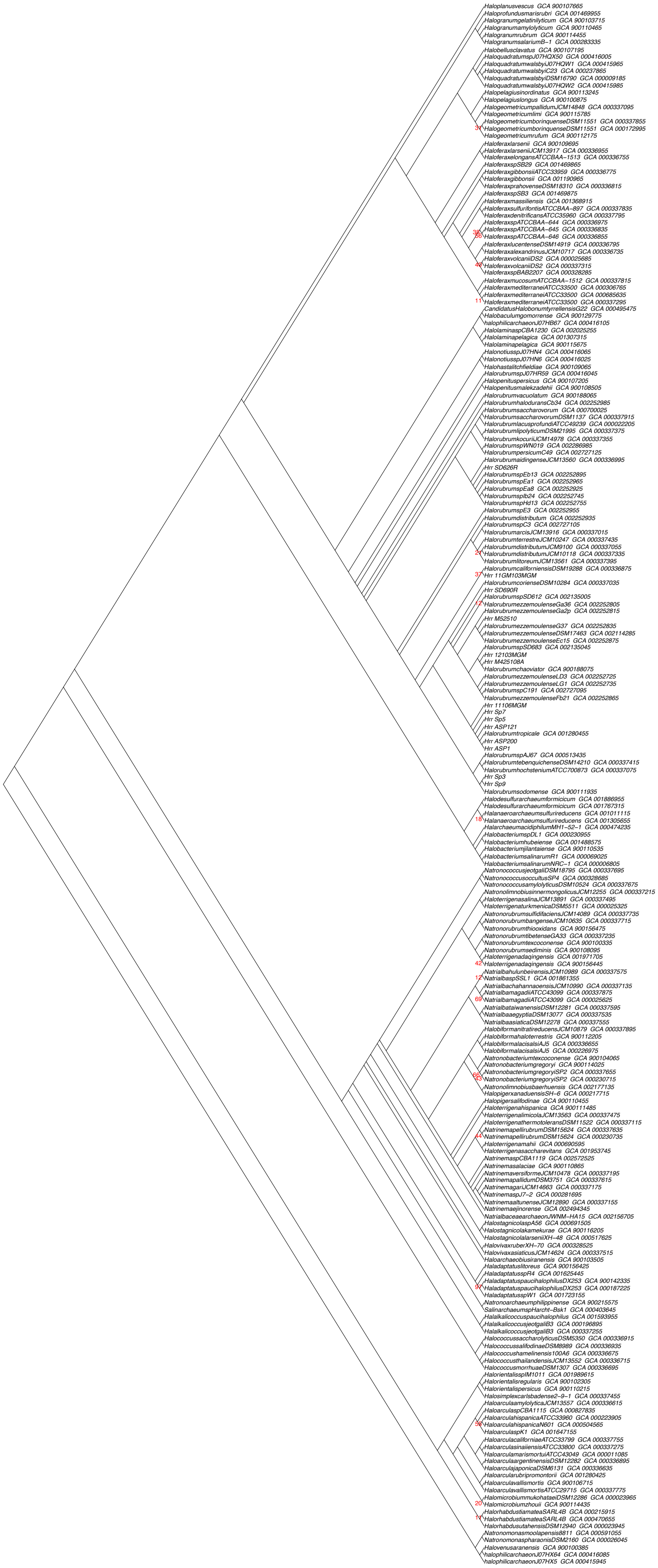
Haloplanusvescus GCA 900107665
Haloprofundusmarisrubri GCA 001469955
Halogranumgelatinilyticum GCA 900103715
Halogranumamylolyticum GCA 900110465
Halogranumrubrum GCA 900114455
HalogranumsalariumB−1 GCA 000283335
Halobellusclavatus GCA 900107195
HaloquadratumspJ07HQX50 GCA 000416005
HaloquadratumwalsbyiJ07HQW1 GCA 000415965
HaloquadratumwalsbyiC23 GCA 000237865
HaloquadratumwalsbyiDSM16790 GCA 000009185
HaloquadratumwalsbyiJ07HQW2 GCA 000415985
Halopelagiusinordinatus GCA 900113245
Halopelagiuslongus GCA 900100875
HalogeometricumpallidumJCM14848 GCA 000337095
Halogeometricumlimi GCA 900115785
HalogeometricumborinquenseDSM11551 GCA 000337855
31
HalogeometricumborinquenseDSM11551 GCA 000172995
Halogeometricumrufum GCA 900112175
Haloferaxlarsenii GCA 900109695
HaloferaxlarseniiJCM13917 GCA 000336955
HaloferaxelongansATCCBAA−1513 GCA 000336755
HaloferaxspSB29 GCA 001469865
HaloferaxgibbonsiiATCC33959 GCA 000336775
Haloferaxgibbonsii GCA 001190965
HaloferaxprahovenseDSM18310 GCA 000336815
HaloferaxspSB3 GCA 001469875
Haloferaxmassiliensis GCA 001368915
HaloferaxsulfurifontisATCCBAA−897 GCA 000337835
HaloferaxdenitrificansATCC35960 GCA 000337795
HaloferaxspATCCBAA−644 GCA 000336975
38
HaloferaxspATCCBAA−645 GCA 000336835
56
HaloferaxspATCCBAA−646 GCA 000336855
HaloferaxlucentenseDSM14919 GCA 000336795
HaloferaxalexandrinusJCM10717 GCA 000336735
HaloferaxvolcaniiDS2 GCA 000025685
40
HaloferaxvolcaniiDS2 GCA 000337315
HaloferaxspBAB2207 GCA 000328285
HaloferaxmucosumATCCBAA−1512 GCA 000337815
HaloferaxmediterraneiATCC33500 GCA 000306765
HaloferaxmediterraneiATCC33500 GCA 000685635
11
HaloferaxmediterraneiATCC33500 GCA 000337295
CandidatusHalobonumtyrrellensisG22 GCA 000495475
Halobaculumgomorrense GCA 900129775
halophilicarchaeonJ07HB67 GCA 000416105
HalolaminaspCBA1230 GCA 002025255
Halolaminapelagica GCA 001307315
Halolaminapelagica GCA 900115675
HalonotiusspJ07HN4 GCA 000416065
HalonotiusspJ07HN6 GCA 000416025
Halohastalitchfieldiae GCA 900109065
HalorubrumspJ07HR59 GCA 000416045
Halopenituspersicus GCA 900107205
Halopenitusmalekzadehii GCA 900108505
Halorubrumvacuolatum GCA 900188065
HalorubrumhaloduransCb34 GCA 002252985
Halorubrumsaccharovorum GCA 000700025
HalorubrumsaccharovorumDSM1137 GCA 000337915
HalorubrumlacusprofundiATCC49239 GCA 000022205
HalorubrumlipolyticumDSM21995 GCA 000337375
HalorubrumkocuriiJCM14978 GCA 000337355
HalorubrumspWN019 GCA 002286985
HalorubrumpersicumC49 GCA 002727125
HalorubrumaidingenseJCM13560 GCA 000336995
Hrr SD626R
HalorubrumspEb13 GCA 002252895
HalorubrumspEa1 GCA 002252965
HalorubrumspEa8 GCA 002252925
HalorubrumspIb24 GCA 002252745
HalorubrumspHd13 GCA 002252755
HalorubrumspE3 GCA 002252955
Halorubrumdistributum GCA 002252935
HalorubrumspC3 GCA 002727105
HalorubrumarcisJCM13916 GCA 000337015
HalorubrumterrestreJCM10247 GCA 000337435
HalorubrumdistributumJCM9100 GCA 000337055
21
HalorubrumdistributumJCM10118 GCA 000337335
HalorubrumlitoreumJCM13561 GCA 000337395
HalorubrumcaliforniensisDSM19288 GCA 000336875
37
Hrr 11GM103MGM
HalorubrumcorienseDSM10284 GCA 000337035
Hrr SD690R
HalorubrumspSD612 GCA 002135005
12
HalorubrumezzemoulenseGa36 GCA 002252805
HalorubrumezzemoulenseGa2p GCA 002252815
Hrr M52510
HalorubrumezzemoulenseG37 GCA 002252835
HalorubrumezzemoulenseDSM17463 GCA 002114285
HalorubrumezzemoulenseEc15 GCA 002252875
HalorubrumspSD683 GCA 002135045
Hrr 12103MGM
Hrr M425108A
Halorubrumchaoviator GCA 900188075
HalorubrumezzemoulenseLD3 GCA 002252725
HalorubrumezzemoulenseLG1 GCA 002252735
HalorubrumspC191 GCA 002727095
HalorubrumezzemoulenseFb21 GCA 002252865
Hrr 11106MGM
Hrr Sp7
Hrr Sp5
Hrr ASP121
Halorubrumtropicale GCA 001280455
Hrr ASP200
Hrr ASP1
HalorubrumspAJ67 GCA 000513435
HalorubrumtebenquichenseDSM14210 GCA 000337415
HalorubrumhochsteniumATCC700873 GCA 000337075
Hrr Sp3
Hrr Sp9
Halorubrumsodomense GCA 900111935
Halodesulfurarchaeumformicicum GCA 001886955
Halodesulfurarchaeumformicicum GCA 001767315
Halanaeroarchaeumsulfurireducens GCA 001011115
18
Halanaeroarchaeumsulfurireducens GCA 001305655
HalarchaeumacidiphilumMH1−52−1 GCA 000474235
HalobacteriumspDL1 GCA 000230955
Halobacteriumhubeiense GCA 001488575
Halobacteriumjilantaiense GCA 900110535
HalobacteriumsalinarumR1 GCA 000069025
HalobacteriumsalinarumNRC−1 GCA 000006805
NatronococcusjeotgaliDSM18795 GCA 000337695
NatronococcusoccultusSP4 GCA 000328685
NatronococcusamylolyticusDSM10524 GCA 000337675
NatronolimnobiusinnermongolicusJCM12255 GCA 000337215
HaloterrigenasalinaJCM13891 GCA 000337495
HaloterrigenaturkmenicaDSM5511 GCA 000025325
NatronorubrumsulfidifaciensJCM14089 GCA 000337735
NatronorubrumbangenseJCM10635 GCA 000337715
Natronorubrumthiooxidans GCA 900156475
NatronorubrumtibetenseGA33 GCA 000337235
Natronorubrumtexcoconense GCA 900100335
Natronorubrumsediminis GCA 900108095
Haloterrigenadaqingensis GCA 001971705
42
Haloterrigenadaqingensis GCA 900156445
NatrialbahulunbeirensisJCM10989 GCA 000337575
12
NatrialbaspSSL1 GCA 001861355
NatrialbachahannaoensisJCM10990 GCA 000337135
NatrialbamagadiiATCC43099 GCA 000337875
69
NatrialbamagadiiATCC43099 GCA 000025625
NatrialbataiwanensisDSM12281 GCA 000337595
NatrialbaaegyptiaDSM13077 GCA 000337535
NatrialbaasiaticaDSM12278 GCA 000337555
HalobiformanitratireducensJCM10879 GCA 000337895
Halobiformahaloterrestris GCA 900112205
HalobiformalacisalsiAJ5 GCA 000336655
HalobiformalacisalsiAJ5 GCA 000226975
Natronobacteriumtexcoconense GCA 900104065
Natronobacteriumgregoryi GCA 900114025
NatronobacteriumgregoryiSP2 GCA 000337655
66
43
NatronobacteriumgregoryiSP2 GCA 000230715
Natronolimnobiusbaerhuensis GCA 002177135
HalopigerxanaduensisSH−6 GCA 000217715
Halopigersalifodinae GCA 900110455
Haloterrigenahispanica GCA 900111485
HaloterrigenalimicolaJCM13563 GCA 000337475
HaloterrigenathermotoleransDSM11522 GCA 000337115
NatrinemapellirubrumDSM15624 GCA 000337635
44
NatrinemapellirubrumDSM15624 GCA 000230735
Haloterrigenamahii GCA 000690595
Haloterrigenasaccharevitans GCA 001953745
NatrinemaspCBA1119 GCA 002572525
Natrinemasalaciae GCA 900110865
NatrinemaversiformeJCM10478 GCA 000337195
NatrinemapallidumDSM3751 GCA 000337615
NatrinemagariJCM14663 GCA 000337175
NatrinemaspJ7−2 GCA 000281695
NatrinemaaltunenseJCM12890 GCA 000337155
Natrinemaejinorense GCA 002494345
NatrialbaceaearchaeonJWNM−HA15 GCA 002156705
HalostagnicolaspA56 GCA 000691505
Halostagnicolakamekurae GCA 900116205
HalostagnicolalarseniiXH−48 GCA 000517625
HalovivaxruberXH−70 GCA 000328525
HalovivaxasiaticusJCM14624 GCA 000337515
Haloarchaeobiusiranensis GCA 900103505
Haladaptatuslitoreus GCA 900156425
HaladaptatusspR4 GCA 001625445
HaladaptatuspaucihalophilusDX253 GCA 900142335
97
HaladaptatuspaucihalophilusDX253 GCA 000187225
HaladaptatusspW1 GCA 001723155
Natronoarchaeumphilippinense GCA 900215575
SalinarchaeumspHarcht−Bsk1 GCA 000403645
Halalkalicoccuspaucihalophilus GCA 001593955
HalalkalicoccusjeotgaliB3 GCA 000196895
HalalkalicoccusjeotgaliB3 GCA 000337255
HalococcussaccharolyticusDSM5350 GCA 000336915
HalococcussalifodinaeDSM8989 GCA 000336935
Halococcushamelinensis100A6 GCA 000336675
HalococcusthailandensisJCM13552 GCA 000336715
HalococcusmorrhuaeDSM1307 GCA 000336695
HalorientalisspIM1011 GCA 001989615
Halorientalisregularis GCA 900102305
Halorientalispersicus GCA 900110215
Halosimplexcarlsbadense2−9−1 GCA 000337455
HaloarculaamylolyticaJCM13557 GCA 000336615
HaloarculaspCBA1115 GCA 000827835
HaloarculahispanicaATCC33960 GCA 000223905
55
HaloarculahispanicaN601 GCA 000504565
HaloarculaspK1 GCA 001647155
HaloarculacaliforniaeATCC33799 GCA 000337755
HaloarculasinaiiensisATCC33800 GCA 000337275
HaloarculamarismortuiATCC43049 GCA 000011085
HaloarculaargentinensisDSM12282 GCA 000336895
HaloarculajaponicaDSM6131 GCA 000336635
Haloarcularubripromontorii GCA 001280425
Haloarculavallismortis GCA 900106715
HaloarculavallismortisATCC29715 GCA 000337775
HalomicrobiummukohataeiDSM12286 GCA 000023965
20
Halomicrobiumzhouii GCA 900114435
HalorhabdustiamateaSARL4B GCA 000215915
11
HalorhabdustiamateaSARL4B GCA 000470655
HalorhabdusutahensisDSM12940 GCA 000023945
Natronomonasmoolapensis8811 GCA 000591055
NatronomonaspharaonisDSM2160 GCA 000026045
Halovenusaranensis GCA 900100385
halophilicarchaeonJ07HX64 GCA 000416085
halophilicarchaeonJ07HX5 GCA 000415945

**Figure S5.** Bootstrap support values of the presence-absence phylogeny mapped onto the ribosomal protein reference tree. All nodes with values of 10% or greater are shown. Only few groupings of organisms belonging to the same species or genus are moderately supported by both, the presence-absence phylogeny and the ribosomal proteins. The lack of mutual support between ribosomal reference tree and the RM system derived phylogeny confirms the discord between the datasets (ribosomal proteins and RM system related proteins).
